# Repeated intraperitoneal administration of the GABA_B_ receptor agonist baclofen reduces body weight in the mouse

**DOI:** 10.1101/2025.01.05.631370

**Authors:** Ivor S Ebenezer

**Affiliations:** Neuropharmacologist Research Group, School of Medicine, Pharmacy and Biomedical Sciences, St Michael’s Building, University of Portsmouth, PO1 2DT, United Kingdom

**Keywords:** GABA, GABA_B_ receptors, Baclofen, Mouse, Food Intake, Body Weight

## Abstract

Chronic intraperitoneal (ip) administration of the GABA_B_ receptor agonist baclofen has been shown to reduce body weight in rats without significantly affecting daily food intake. The present study was undertaken to extend these observations to another rodent species. Male C57B/10 mice (n=27) that had free access to food and water were injected once daily for 15 days with either saline or baclofen (4 or 8 mg / kg; i.p). Body weight and food intake were measured 24h after each injection. Statistical analysis of the results revealed that the 4 mg / kg dose of baclofen significantly reduced body weight gain in the mice during the first 5 days while the 8 mg / kg dose reduced body weight gain throughout the 15 days of the study when compared with saline control animals (F(_2,21_)=4.88, P<0.02). Baclofen (4 or 8 mg / kg) generally had no significant effects on daily 24h food intake except on days 1 and 2 when the 8 mg / kg dose produced small but significant decreases in food intake (P<0.05) that were probably due to the initial depressant effects of the drug on behaviour. The results of this study extend previous findings in rat to another rodent species and show that systemic administration of the GABA_B_ receptor agonist baclofen reduces body weight gain in the mouse without affecting daily food intake. These results lend further support to the hypothesis that GABA_B_ receptor agonists decreases body weight by increasing metabolic rate.

## 1. Introduction

It has been well established that intracerebroventricular (icv) or systemic injections of the GABA_B_ receptor agonist baclofen (see Ebenezer, 2015) increases food intake in satiated or free feeding animals including pig, rat and mouse by an action at central GABA_B_ receptors (Ebenezer, 1990, Ebenezer and Baldwin, 1990, Ebenezer and Pringle, 1992, Ebenezer, 1995, Ebenezer, 2012, Ebenezer and Patel, 2004, Higgs and Barber, 2004, Ebenezer and Patel, 2004, Buda-Levin et al., 2005, Patel Ebenezer, 2008a and Ebenezer and Prabhaker, 2007). Furthermore, we have demonstrated that endogenous GABA, acting at central GABA_B_ receptors plays a physiological role in the regulation of feeding behaviour (Patel and Ebenezer, 2004). Pharmacological and immunohistochemical studies have implicated a number of brain areas, such as the medial raphe nucleus, the nucleus accumbens and the arcuate nucleus of the hypothalamus, as possible central sites where the GABA or GABA_B_ receptor agonists act to mediate their hyperphagic actions (Wirtshafter et al., 1993, Stratford and Kelly, 1997, Ward et al., 2000, Backberg et al., 2003).

While most of the studies have focussed primarily on the effects of acute administration of the GABA_B_ receptor agonist on short-term feeding responses (Ebenezer, 1990, Ebenezer, 1995, Ebenezer, 1996, Ebenezer and Pringle, 1992, Ebenezer and Patel, 2004, Patel and Ebenezer, 2008b, Ebenezer and Prabhaker, 2007, Stratford and Kelly, 1997, Ward et al., 2000, Higgs and Barber, 2004), we have also investigated the effects of chronic injections of baclofen (1 and 4 mg/kg, administered i.p. once daily), on short- and long term (24h) food intake and body weight in non-deprived rats (Patel and Ebenezer, 2010). We found that (i) baclofen (1 and 4 mg/kg., i.p.) elicits a short-term (30 min) hyperphagia in the animals over the 27 day treatment period, (ii) tolerance did not occur to the short-term hyperphagia over the treatment period, (iii) long term (24 h) food intake following acute and chronic exposure to baclofen (1 and 4 mg/kg, i.p.) was not significantly different from the controls, suggesting that the animals were able to regulate their daily intake quite accurately, and (iv) chronic administration of baclofen (4 mg/kg but not the 1 mg/kg dose) significantly reduces body weight. The observations that chronic administration of baclofen (4 mg/kg) stimulates short-term food intake without affecting long term (24 h) feeding but decreases body weight suggest that baclofen may act through different mechanisms to influence food intake and body weight.

The present study was undertaken to extend these findings to another rodent species by investigating the effects of repeated administration of baclofen on daily food consumption and body weight in the mouse.

## 2. Material and methods

The protocols used in this study were approved by the Ethical Review Committee at the University of Portsmouth, England and carried out under licences granted by the UK Home Office Scientific Procedures Act.

### 2.1 Effects of repeated administration of baclofen on food intake and body weight

Adult male C57B/10 mice (n=23; starting body weights: 20 - 28.6 g) were housed in cages dimensions: 33×20×18 cm) in groups of 2 (except 1 mouse that was housed singly) where they had free access to food (Food composition: protein 20%, oil 4.5%, carbohydrate 60%, fibre 5%, ash, 7% + traces of vitamins and metals) and water at all times. Prior to the start of the experiment, the animals were handled and their body weights measured for 10 days to get them used the experimental procedure. The food was presented to the mice in a food hopper that could be removed and the food tipped into a plastic beaker so that it could be weighed. During the experimental sessions that followed, the mice were injected i.p. with either physiological saline solution (n=8), baclofen (4 mg/kg; n=7) or baclofen (8 mg/kg; n=8) at between 14.00 - 14.30 each day for 15 days. The weights of each of the mice were recorded prior to injection. In addition, the daily amount of food consumed by the mice in each cage was calculated by placing a known weight of food in the food hopper and measuring the weight of the food 24 later.

### 2.2 Drugs

(±) Baclofen was purchased from Sigma Biochemicals, Dorset, UK. The drug was dissolved in physiological saline solution (0.9% w/v, NaCl) to give an injection volume of 0.1 ml / 100 g body weight. Physiological saline solution was used in control experiments.

### 2.3 Statistics

The data from was analysed by 2 way analysis of variance (ANOVA) with repeated measures on treatment and time (days) followed by the Student-Newman Keul *post-hoc* test (**Winer, 1971**).

### 2.4 Food Intake and Body Weight

The food intake data was expressed as g per 100 g body weight of the animals in each cage.

The body weight data obtained for each mouse were expressed as a percentage of the average animal’s body weight recorded on the day prior to and the day the animals received their first injections.

## 3. Results

### 3.1. Effects of repeated administration of baclofen on food intake and body weight in free feeding mice

#### 3.1.1. Food Intake

The effects of baclofen (4 or 8 mg/kg, i.p.) or physiological saline on the amount of food consumed by the mice in each cage recorded daily over the 15 day recording period are illustrated in Fig. 1. Statistical analysis of the data showed that there were significant main effects of treatment (F*(2,9)* = 6.3, *P* < 0.05) and time (days) (F*(14,126)* = 2.56, *P* < 0.01), but no significant effects of treatment x time (days) interaction (*F*(28,126)= 1.47, *NS*). *Post-hoc* tests revealed that repeated treatment with baclofen 8 mg/kg produced small but significant decreases (P<0.05) compared with saline controls in food intake on Days 1 and 2 only Fig. 1B). There were no significant effects on food consumption between the saline and baclofen 4mg / kg groups on Days 1 to 15 (Fig. 1A).

**Figure 1.**
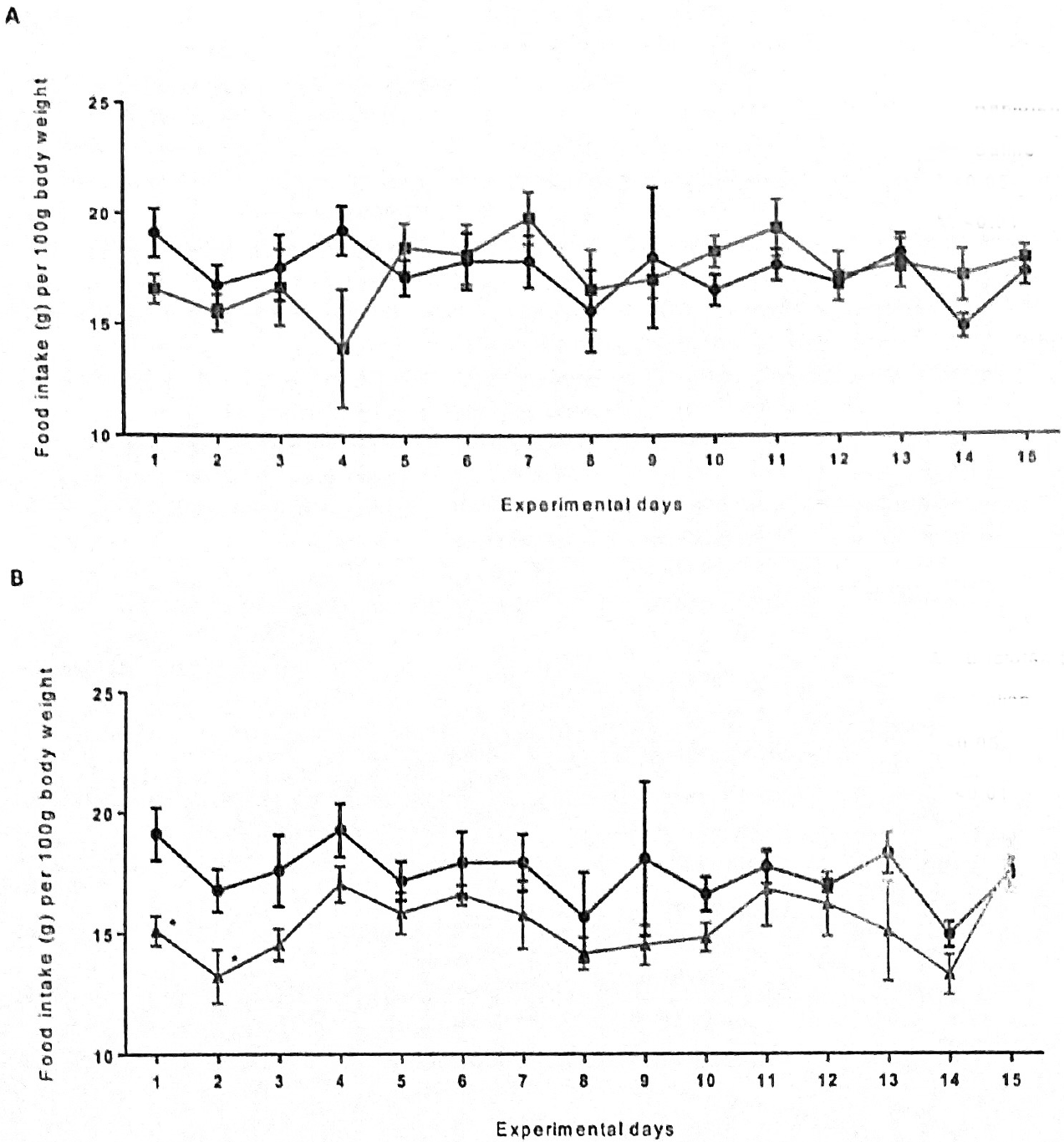
Effects of repeated once daily intraperitoneal injections of physiological saline (●) or (A) 4 mg/kg baclofen 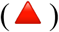 and (B) 8 mg/kg baclofen (◼) on daily 24h food intake in mice expressed as g per 100 g body weight recorded over a period of 15 days (body weight = 20 - 28 g at start of experiment). See text for further details and statistical analyses. Vertical lines represent ± S.E.M.

#### 3.1.3. Body Weight

The effects of baclofen (4 or 8 mg/kg, i.p.) or physiological saline on body weight after each feeding session during Days 1 to 15 are shown in Fig. 2. Statistical analysis of the results revealed that there were significant main effects of treatment (F*(2,21)* = 4.88, *P* < 0.02) and time (days) (F*(14,294)* = 36.7, *P* < 0.01), and significant effects of treatment x time (days) interaction (*F*(28,294) = 46.0, *P* < 0.01). *Post-hoc* tests showed that the 4mg / kg dose of baclofen produced significant reduction in body weight gain compared to controls after the first 5 days of treatment (P<0.05 for each day, Fig. 2A). However, the 8mg/kg dose significantly decreased body weight gain compared with controls throughout the 15 days of the experiment (at least P<0.05 for each day, Fig. 2B).

**Figure 2.**
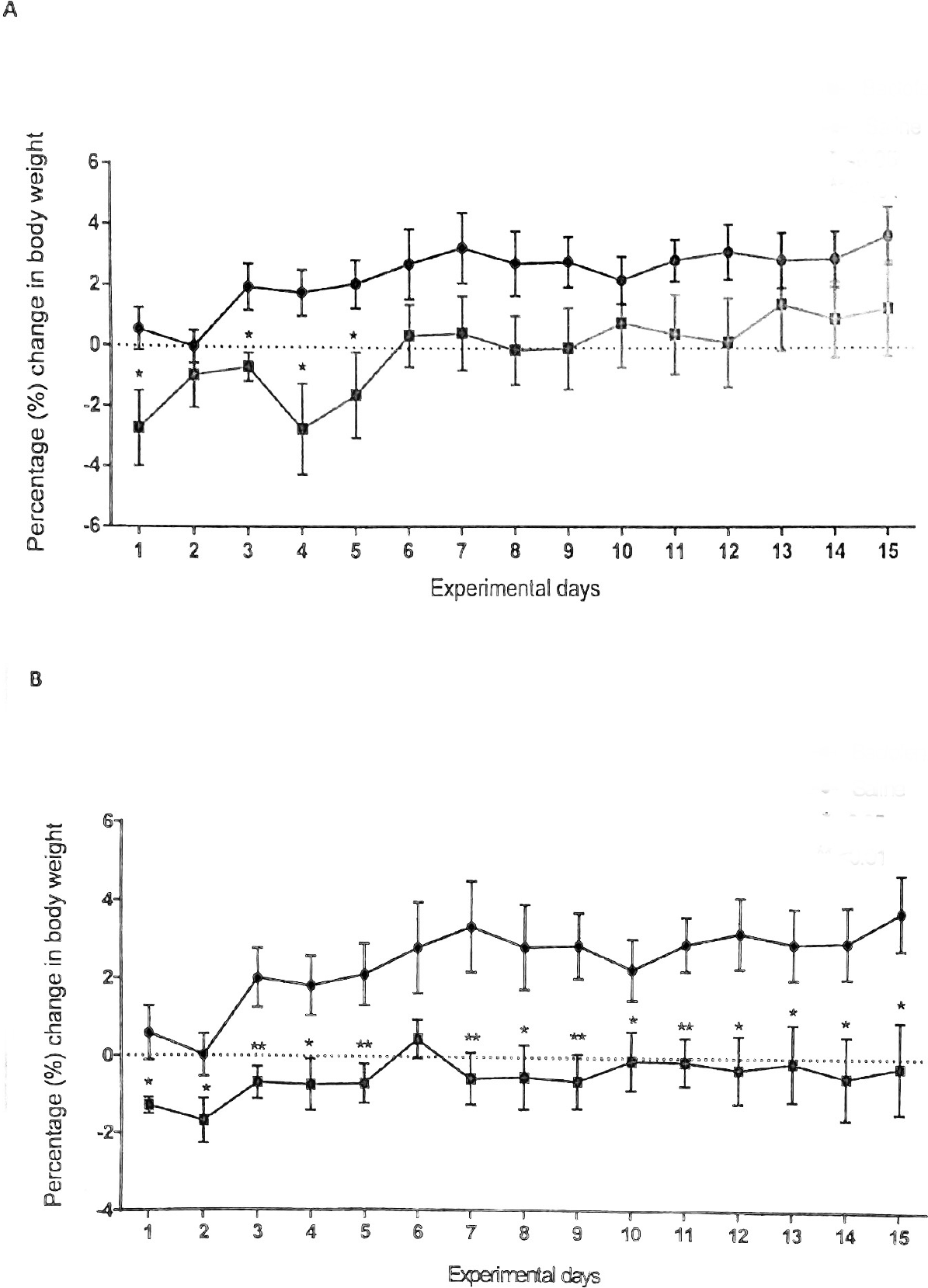
Effects of repeated once daily intraperitoneal injections of physiological saline (●) or (A) 4 mg/kg baclofen (◼) and (B) 8 mg/kg baclofen (◼) on percentage change in body weight in mice recorded over a period of 15 days (body weight = 20 - 28 g at start of experiment). See text for further details and statistical analyses. Vertical lines represent ± S.E.M.

## 4. Discussion

It has previously been reported that repeated systemic administration of baclofen once daily to free feeding rats reduces body weight without affecting 24h food intake (**Patel and Ebenezer, 2010**). The present study was undertaken to extend these observations by investigating the effects of repeated administration of baclofen on daily food intake and body weight in another rodent species.

Doses of baclofen (4 and 8 mg/kg) that have previously been shown to increase food intake in free feeding mice were used in the present study. The results show that both doses of baclofen had little or no effect on daily food consumption but reduced body weight.

The 4 mg/kg dose of baclofen had no effect on daily food intake throughout the study. In contrast, the 8 mg/kg dose caused small but significant decreases in food intake on days 1 and 2. This initial reduction likely resulted from a transient depressant effect of the drug on behaviour, as higher doses of baclofen can induce ataxia, muscle relaxation, and sedation in mice (Ebenezer and Prabhaker, 2007). However, it is likely that the mice rapidly developed tolerance to these behavioural effects (see Ebenezer and Prabhaker, 2007, Patel and Ebenezer, 2010; Bains and Ebenezer, 2013), resulting in no significant differences in daily food intake between the 8 mg/kg group and saline control group from days 3 to 15 (Fig. 1).

The 4 mg/kg baclofen dose significantly reduced body weight gain during the first 5 days of the study, whereas the 8 mg/kg dose reduced body weight gain throughout the 15-day study period (Fig. 2). These findings align with observations in rats, where baclofen-induced weight reduction exhibited a dose-dependent relationship. Previous studies in rats have demonstrated that chronic baclofen administration at lower doses (1 and 2 mg/kg i.p., daily) did not significantly alter body weight (Krolczyk et al., 2005; Patel and Ebenezer, 2008a, Patel and Ebenezer, 2010). However, repeated daily administration of a 4 mg/kg dose produced significant reduction in body weight (Patel and Ebenezer, 2010). Additionally, Rothwell et al. (1985) reported that chronic baclofen administration (2 mg/kg, s.c., twice daily) significantly decreased body weight in rats over 15 days. Notably, they did not assess daily food intake in their study. Considering that baclofen’s half-life in the rat is approximately 4 hours (Popova et al., 1995), a more sustained drug exposure may be required to induce weight reduction. This can be achieved by administering higher doses (e.g., 4 mg/kg once daily; Patel and Ebenezer, 2010) or by more frequent dosing (e.g., 2 mg/kg twice daily; Rothwell et al., 1985). Furthermore, it has been observed that tolerance to the weight-reducing effects of baclofen in rats can develop with repeated administration (Patel, 2010). Therefore, it is plausible that the 4 mg/kg dose in the present study may have been at the lower end of the effective dose range for mice, and the observed reduction in weight gain may have diminished after 5 days due to the development of tolerance. Interestingly, Sato et al. (2007) reported that lean mice chronically exposed to baclofen in their drinking water for 5 weeks did not exhibit any differences in body weight gain compared to vehicle-treated controls. These findings suggest that the blood levels of baclofen achieved through this route may have been insufficient to induce significant weight changes.

The precise mechanisms underlying baclofen’s effects on body weight remain to be fully elucidated. It has been proposed that baclofen may reduce body weight by stimulating thermogenesis in brown adipose tissue (Rothwell et al., 1985, Addae et al., 1986). These authors suggested that this occurs through activation of the sympathetic nervous system, potentially mediated by actions within the ventromedial hypothalamus (VMH). Indeed, Rothwell et al. (1985) demonstrated that baclofen microinjected into the VMH of anaesthetised rats activates brown fat metabolism. However, other mechanisms may also contribute to the weight-reducing effects of chronic baclofen administration. While direct peripheral effects on body weight remain unproven, immunohistochemical studies have revealed that GABA_B_ receptors are associated with neuropeptide Y (NPY) and proopiomelanocortin (POMC) neurones within the arcuate nucleus of the hypothalamus (Backberg et al., 2003). This suggests that baclofen may influence metabolic rate by modulating these hypothalamic neurones, which play crucial roles in regulating energy balance (Morton et al., 2006). Specifically, baclofen could potentially alter the activity of NPY and/or POMC neurones, leading to a decrease in body weight.

In conclusion, this study extends previous observations in rats (Patel and Ebenezer, 2008a, 2010; Ebenezer and Patel, 2011, Ebenezer, 2023) to mice, demonstrating that repeated intraperitoneal injections of baclofen decrease body weight and is not related to suppression of food intake. These findings further support the potential therapeutic use of GABA_B_ agonists, such as baclofen, in the treatment of obesity. This notion is supported by a preliminary open-label clinical trial in obese human subjects, which indicated that baclofen can reduce body weight (Arima and Oiso, 2010).

However, further research is necessary to elucidate the precise mechanisms underlying the effects of baclofen on body weight.

## Acknowledgements

I wish to thank Nasra, Jessica and Kush for assistance in recording body weight and food intake.

